## Supplementary Figures and Tables for "Two subtypes of GTPase-activating proteins coordinate tip growth and cell size regulation in *Physcomitrium patens*"

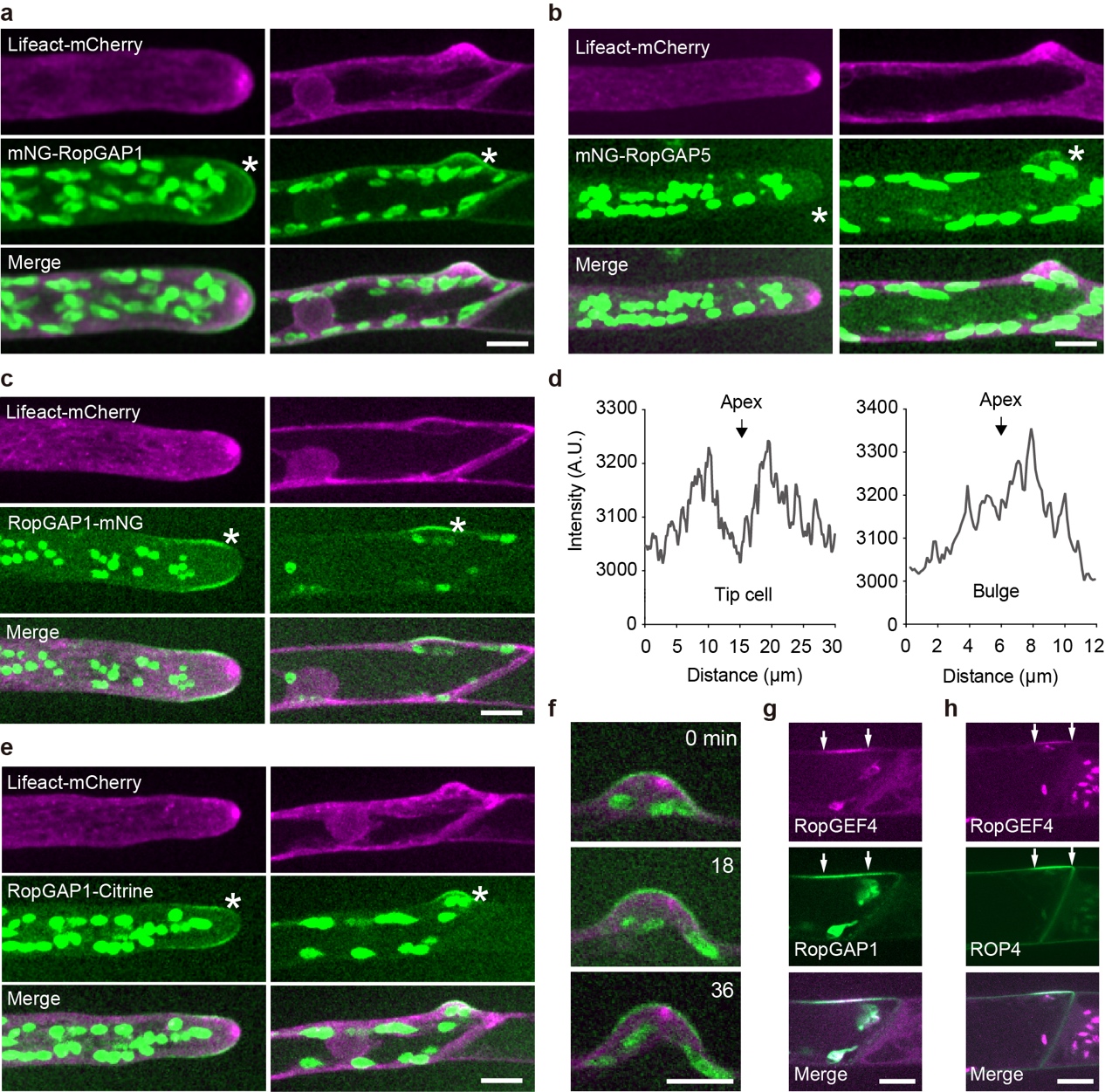

**Supplementary Fig. 1: Localization of PpRopGAP1 in tip cells and subapical branching cells.** (a) Localization of PpRopGAP1 fused with an N-terminal mNG tag. Actin is labeled with Lifeact-mCherry. Stars indicate the apical membrane of a tip cell or the apical membrane of a bulge in the subapical cell. (b) Localization of PpRopGAP5 fused with an N-terminal mNG tag. (c) Localization of PpRopGAP1 fused with a C-terminal mNG tag. (d) Representative intensity plots of PpRopGAP1-mNG along the apical membrane in a tip cell and at the branching site (bulge) in a subapical cell. Note that the low-intensity area is not present at the branching site. (e) Localization of PpRopGAP1 fused with a C-terminal Citrine tag. (f) Time-lapse images of PpRopGAP1-mNG during bulge growth. Note that the actin foci undergo dynamic assembly and disassembly, while PpRopGAP1-mNG does not exhibit a low-intensity zone at the apex. (g) Colocalization of PpRopGAP1-mNG and PpRopGEF4-mCherry before bulge initiation. (h) Colocalization of PpROP4-mNG and PpRopGEF4-mCherry before bulge initiation. Arrows indicate the boundary of PpRopGEF4 fluorescence. Scale bars in all panels: 10 µm.

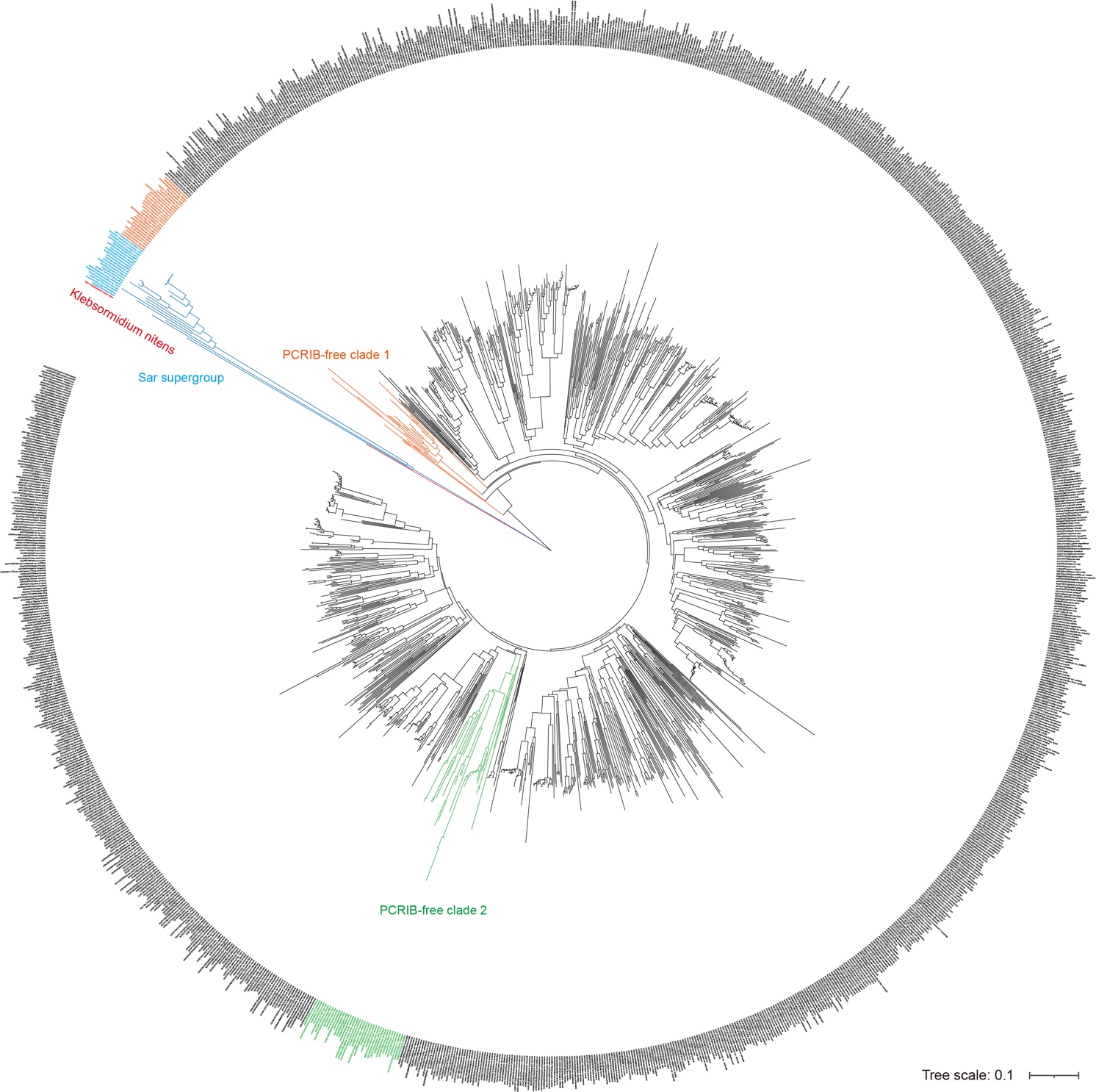

**Supplementary Fig. 2: Phylogenetic tree of RopGAPs.** In total, 1 244 RopGAPs that comprise a CRIB domain and a RhoGAP domain are retrieved from the InterPro database. Protein sequences are obtained from the Uniprot database and aligned by Clustal Omega. The phylogenetic tree is constructed using the neighboring-joining method with a 1 000 bootstrap value in Mega11. The majority of RopGAPs contain a pre-CRIB (PCRIB) motif preceding the CRIB core sequence. The RopGAP homologs in the Sar outgroup and *Klebsormidium nitens* are indicated in blue and red, respectively. Two small subclades that lack the PCRIB motif are shown in orange and green.

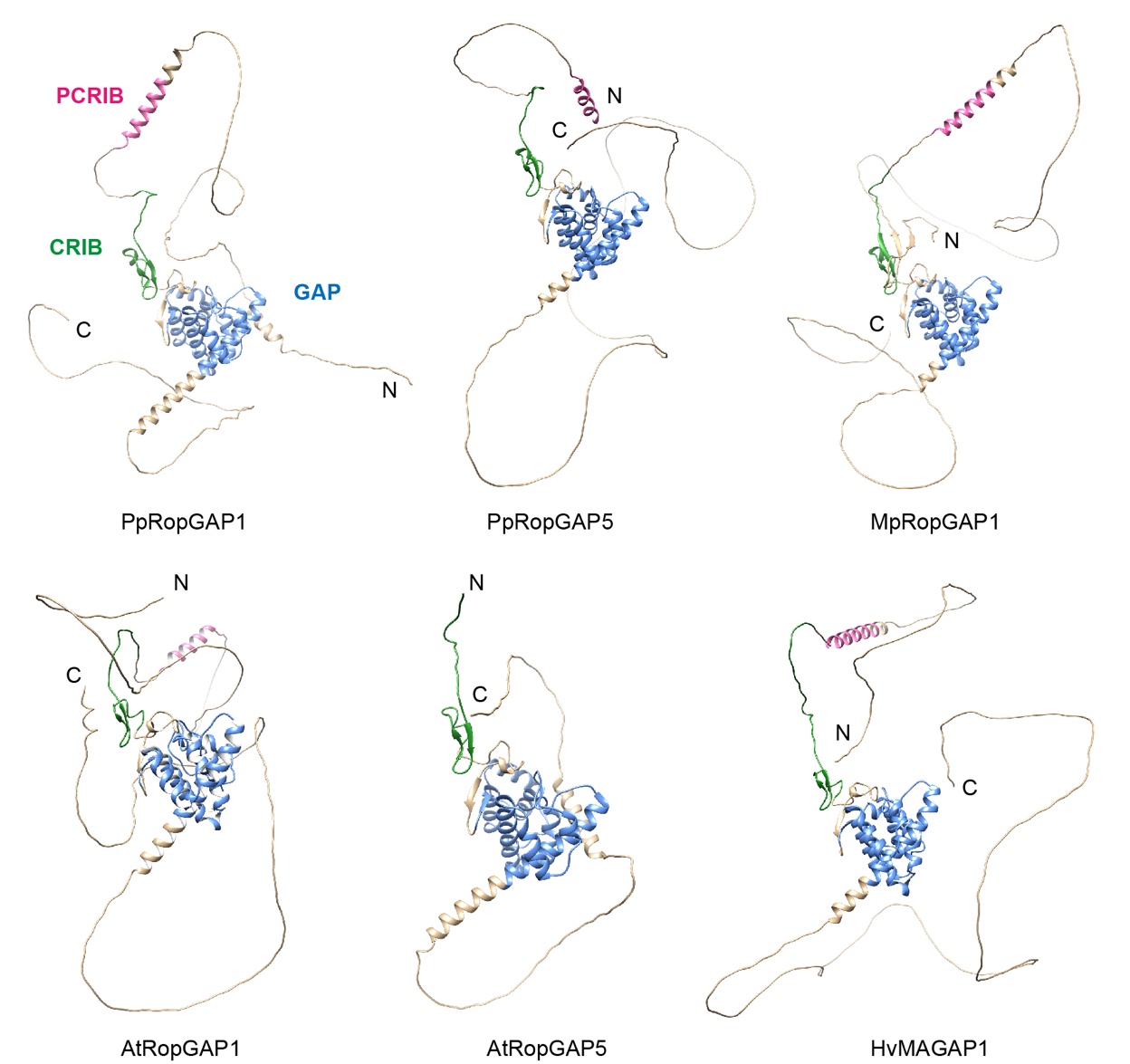

**Supplementary Fig. 3: AlphaFold-predicted structures of RopGAPs.** The N-terminal (N) and C-terminal (C) ends of each protein are indicated. The PCRIB motif, CRIB motif, and GAP domain are shown in magenta, green, and blue, respectively. Note that the PCRIB motif is predicted to fold into an α-helix, but is absent in AtRopGAP5. Pp, *Physcomitrium patens*; Mp, *Marchantia polymorpha*; At, *Arabidopsis thaliana*; Hv, *Hordeum vulgare*.

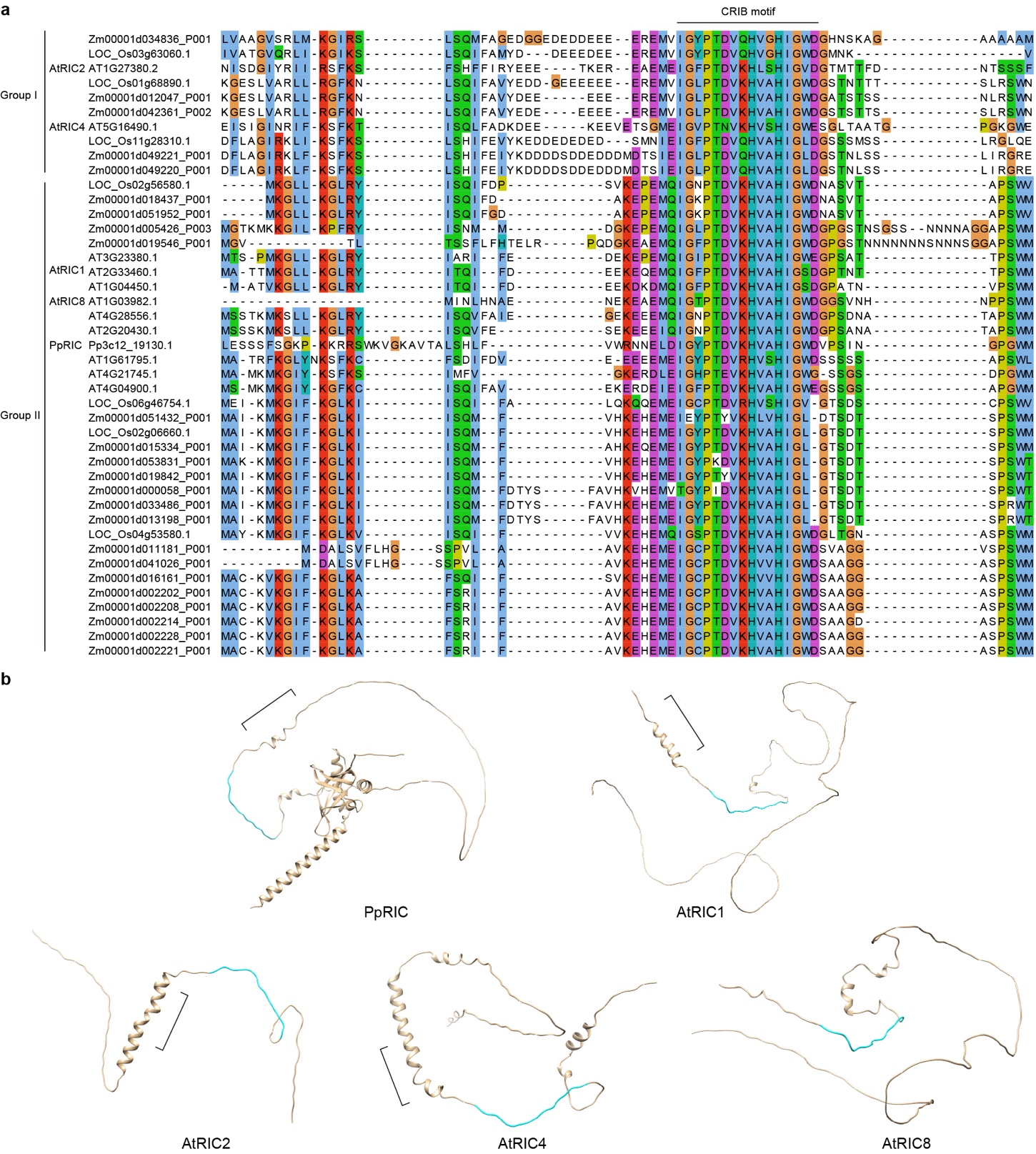

**Supplementary Fig. 4: Sequence alignment and predicted structures of RICs.** (a) Protein sequences of RICs from *Physcomitrium patens*, Arabidopsis, rice, and maize were aligned using Clustal Omega. Phylogenetically, RICs could be classified into two groups. The CRIB motif as well as its flanking regions is shown. Note that the N-terminal and C-terminal regions are conserved in most RICs. Group I RICs contain extra polyacidic residues immediately before the CRIB motif. (b) Alphafold predicted structures of representative RICs. The conserved region (indicated by brackets) before the CRIB motif (cyan) in RICs is different from those in RopGAPs in sequence. However, it also tends to adopt an α-helical structure except for RICs such as AtRIC8 which lack a full-length N-terminal region.

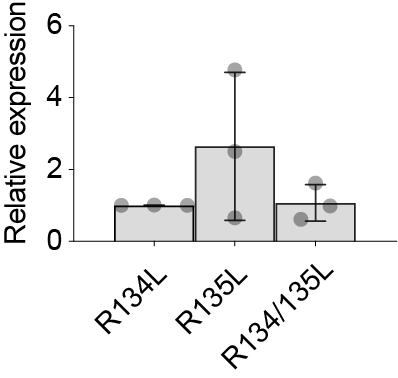

**Supplementary Fig. 5: Relative transcriptional expression levels of PpRopGAP1 with single and double mutations in the PCRIB motif.** The expression levels were detected by qRT-PCR using the TAF10 as the reference gene. Quantification was performed for three independent biological samples from four technical repeats. The expression of PpRopGAP1(R135L) and PpRopGAP1(R134/135L) was normalized to that of PpRopGAP1(R134L). No significant difference was found for each pair of the data using a one-way ANOVA test.

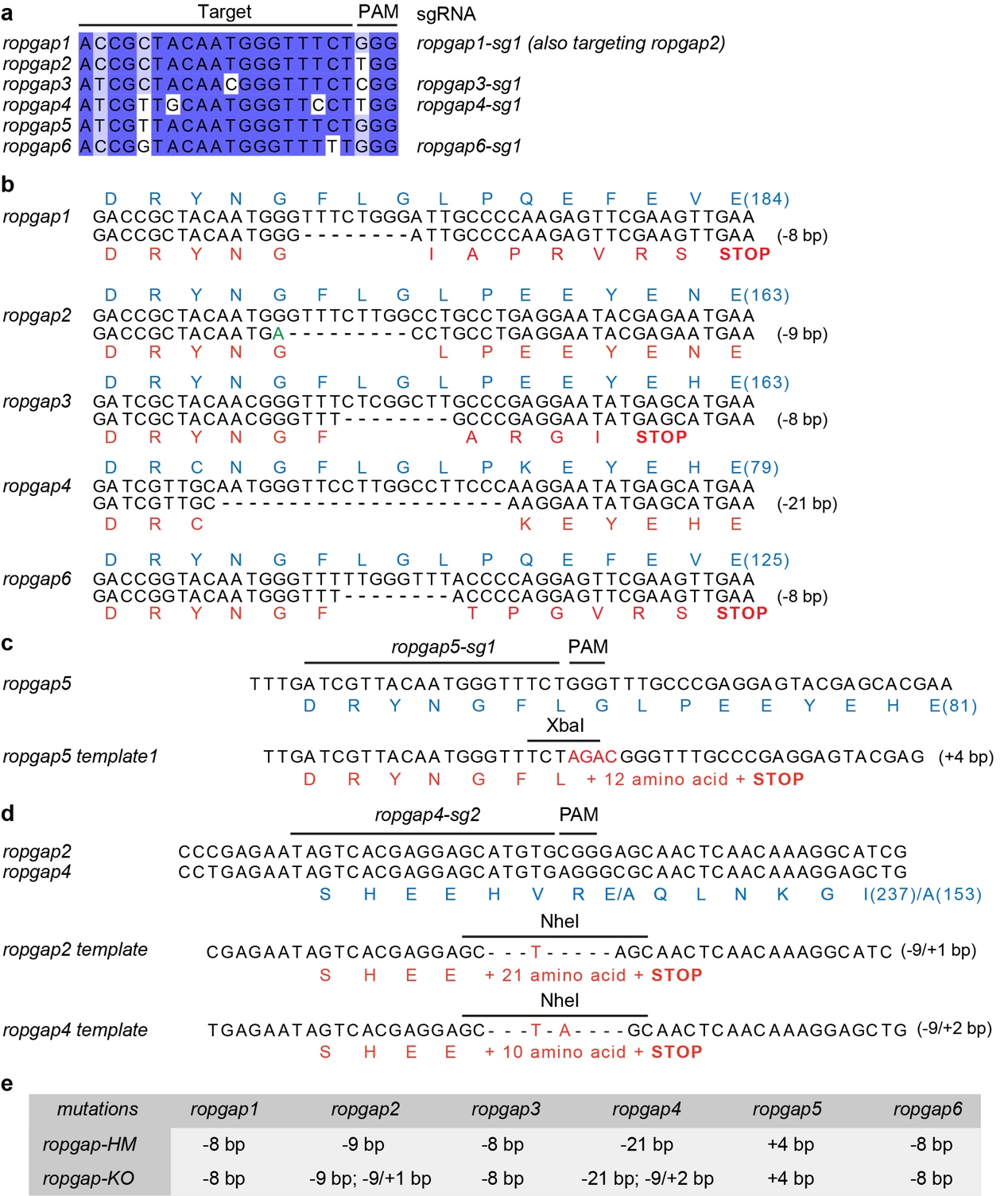

**Supplementary Fig. 6: Construction of the *ropgap* mutants.** (a) Alignment of CRISPR guide RNA (sgRNA) targets used for the first round of genome editing. Four targets (*ropgap1-sg1*, *ropgap3-sg1*, *ropgap4-sg1*, *ropgap6-sg1*) are cotransformed, of which the *ropgap1-sg1* also targets *ropgap2*. PAM, protospacer adjacent motif. (b) Mutations generated by CRISPR/Cas9 editing. The change of translated amino acids (represented by single letters) is shown below the DNA sequences. (c) Oligonucleotide template-dependent genome editing of *ropgap5*. An *Xba* I site is introduced in the template to facilitate genotyping. (d) Oligonucleotide template-dependent genome editing of *ropgap2* and *ropgap4*. The *ropgap4-sg2* is used for simultaneous editing of *ropgap2* and *ropgap4*. A *Nhe* I site is introduced in each template to facilitate genotyping. (e) Summary of mutations in the final *ropgap* sextuple knockout (KO) or hypomorphic (HM) mutants.

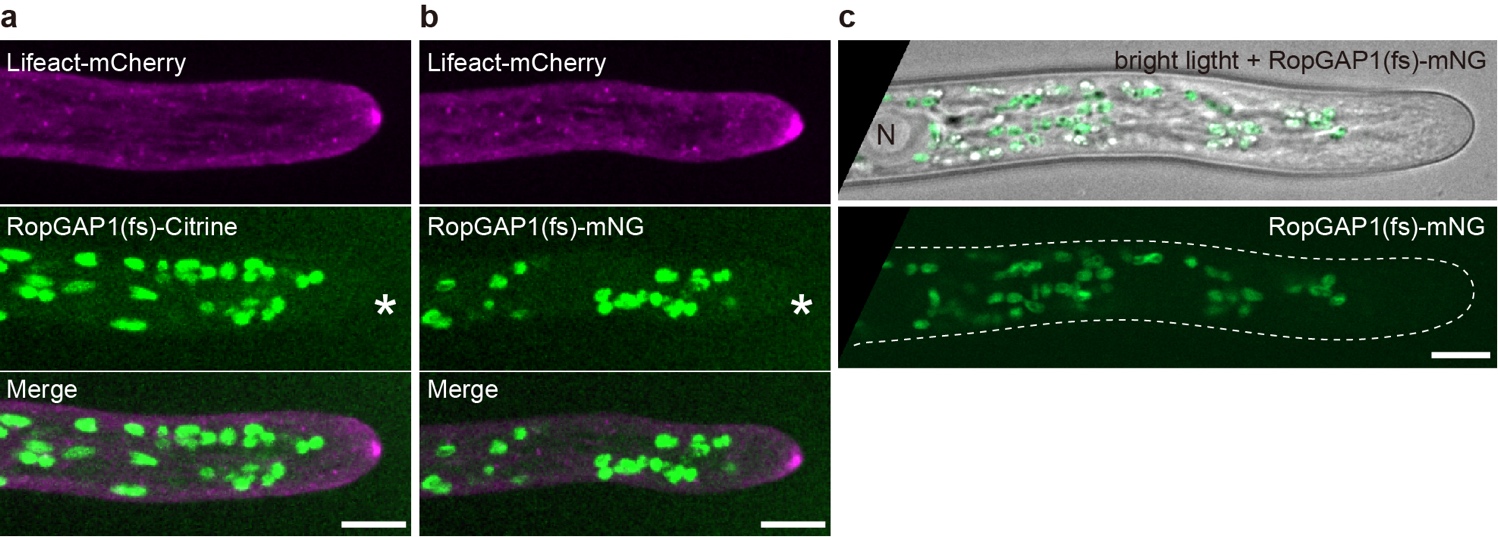

**Supplementary Fig. 7: The expression of PpRopGAP1 is completely blocked by frameshift mutations.** (a) Expression of the endogenous PpRopGAP1 carrying an 8-bp frameshift mutation (fs) in the CRIB domain and fused with a C-terminal Citrine. No fluorescence is detectable at the apical membrane (star). Actin is labeled with a Lifeact-mCherry reporter. (b) Expression of the endogenous PpRopGAP1 carrying an 8-bp frameshift mutation (fs) in the CRIB domain and fused with a C-terminal mNG. (c) Overexpression of the mutated PpRopGAP1 with a C-terminal mNG driven by EF1α promoter. N, nucleus. Scale bars in all panels: 10 µm.

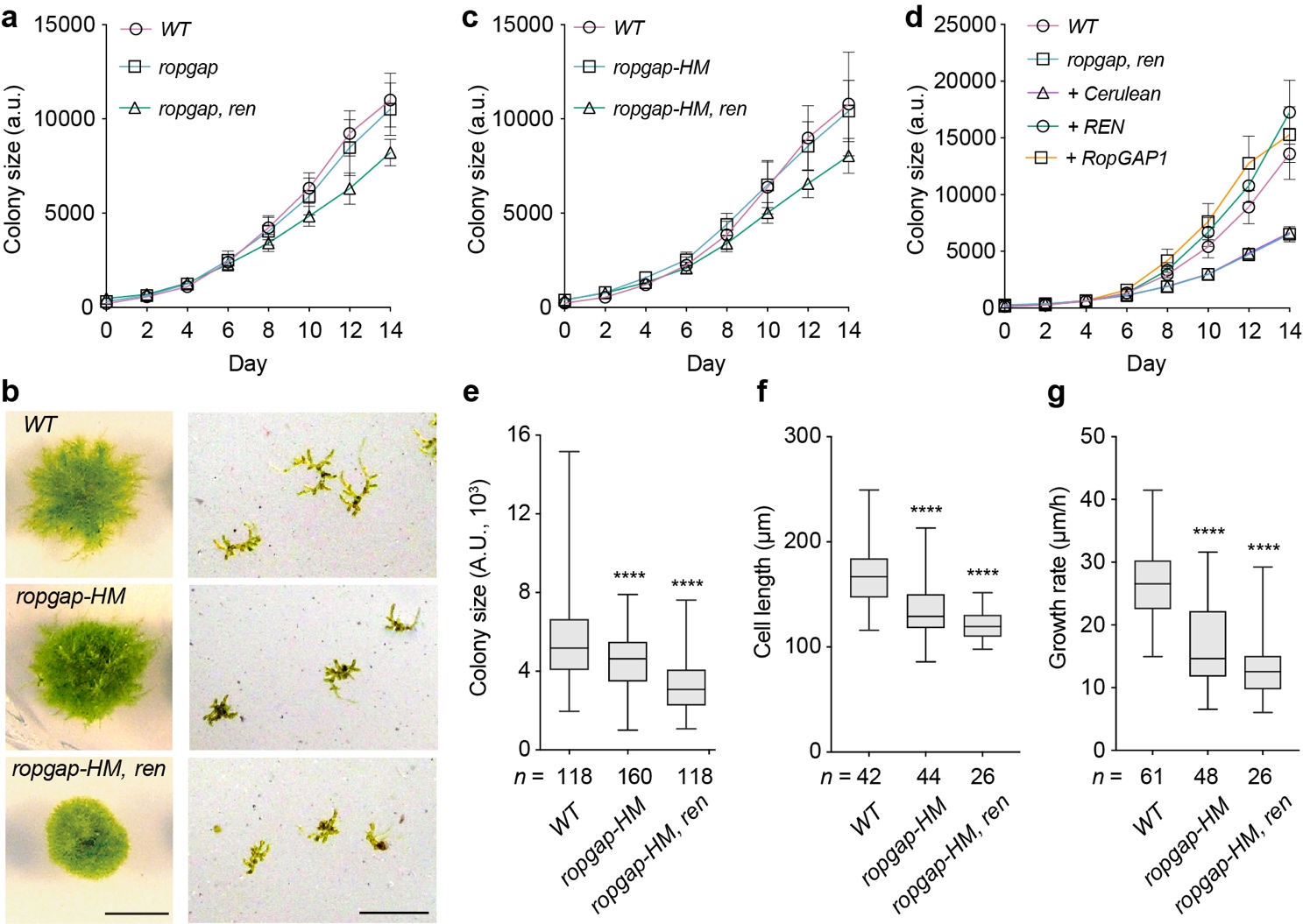

**Supplementary Fig. 8: Phenotypes and rescue of PpRopGAP-related mutants.** (a) Quantification of the colony size of wild-type (WT) and knockout mutant mosses cultured for two weeks. n = 32 colonies (mean ± SD) for each strain. (b) Colony growth and protoplast regeneration in WT, hypomorphic *ropgap* sextuple mutant (*ropgap-HM*), and *ropgap-HM, ren* septuple mutant. Scale bar: 0.5 cm (left), 0.5 mm (right). (c) Quantification of the colony size of WT and hypomorphic mutant mosses cultured for two weeks. n = 32 colonies (mean ± SD) for each strain. (d) Quantification of the colony size of mutant and rescued mosses cultured for two weeks. n = 16 colonies (mean ± SD) for each strain. (e) Quantification of the colony size of 7-day-old regenerated mosses after protoplasting. (f) Quantification of subapical cell length. (g) Quantification of the growth rate of caulonema tip cells. The number of colonies or cells used for quantification is shown at the bottom in (e-g). Each group is compared with WT and analyzed by a one-way ANOVA test. ****, *p* < 0.0001.

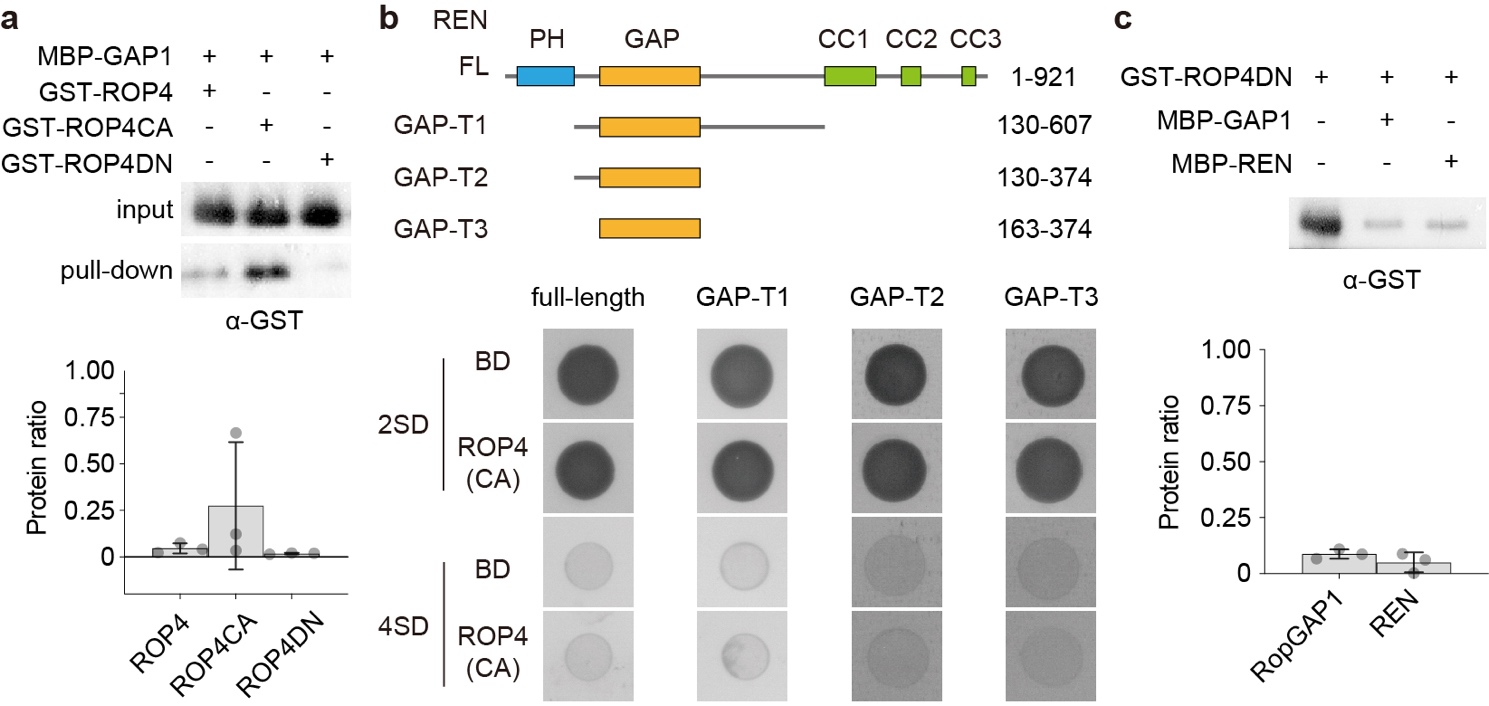

**Supplementary Fig. 9: Analyses of PpRopGAP1 and PpREN interactions with different activity forms of PpROP4.** (a) MBP-PpRopGAP1 preferentially interacts with the active form of PpROP4 (GST-PpROP4CA) as detected by pull-down and western blotting assays. (b) PpREN does not interact with PpROP4CA in yeast-two-hybrid assays. REN’s full-length (FL) and three truncated forms (GAP-T1 to GAP-T3) were tested for interaction with PpROP4CA. PH, PLECKSTRIN HOMOLOGY domain; GAP, GTPASE-ACTIVATING domain; CC, coiled-coil motifs. (c) PpRopGAP1 and PpREN did not exhibit differences in binding capacity with the inactive form of PpROP4 (PpROP4DN). All experiments were performed with at least three replicates.

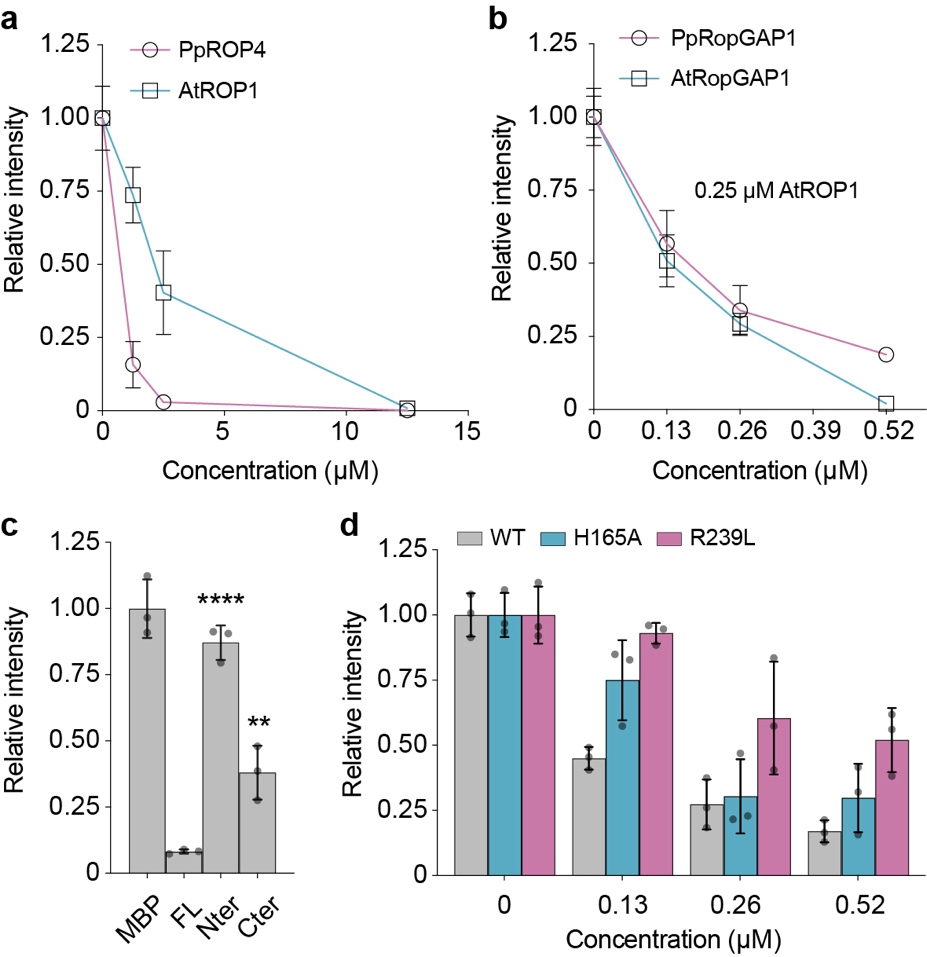

**Supplementary Fig. 10: The GAP activity of wild-type and mutant forms of PpRopGAP1 in vitro.** (a) The intrinsic GTPase activity of PpROP4 and AtROP1. (b) Stimulated GTPase activity of AtROP1 by PpRopGAP1 and AtRopGAP1. (c) The GAP activity of full-length (FL) and truncated forms of PpRopGAP1. Note that the Nter portion of PpRopGAP1 has little GAP activity, while its absence reduces the GAP activity of PpRopGAP1. Each group is compared with the FL PpRopGAP1 using a one-way ANOVA test. **, *p* < 0.01; ****, *p* < 0.0001. (d) The GAP activity of PpRopGAP1 carrying a mutation in the CRIB motif (H165A) or a mutation required for GAP activity (R239L). Note that H165A only reduces GAP activity at low concentrations, indicating that the CRIB motif is not essential for but enhances GAP activity. All data are derived from three replicates and are shown as mean ± SD.

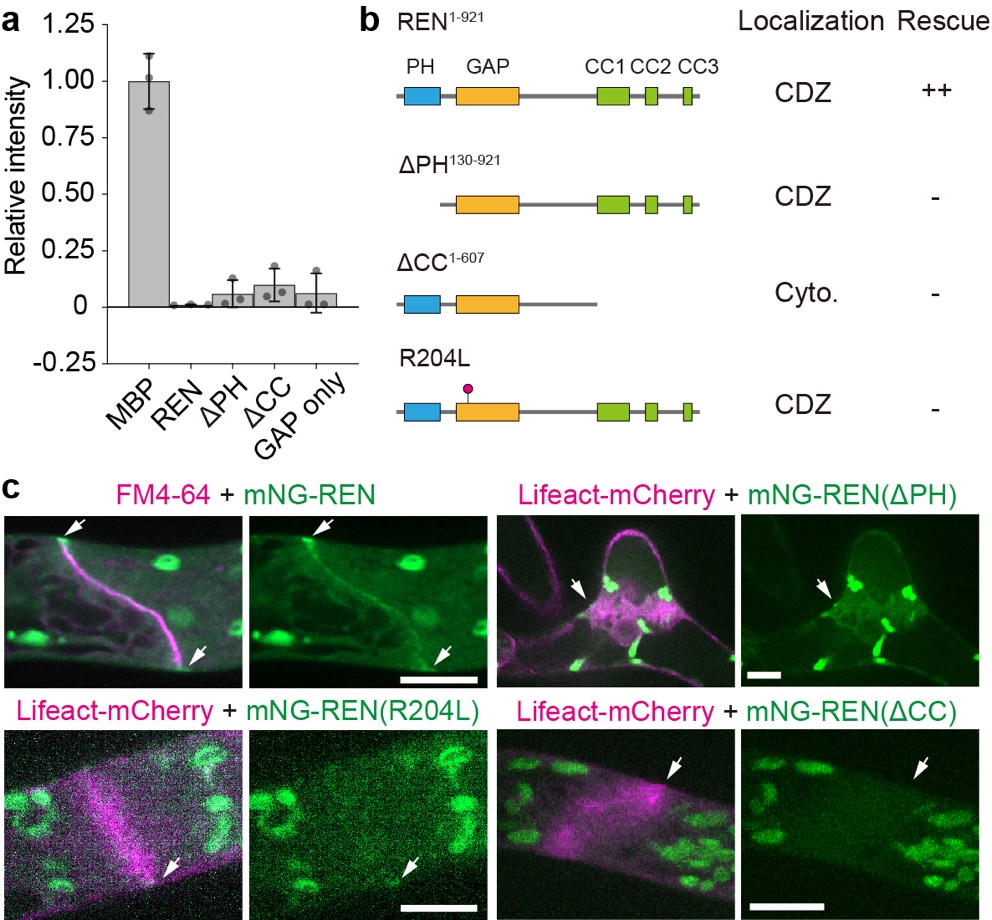

**Supplementary Fig. 11: The functions of PH domain and coiled-coil motif-containing tail of PpREN.** (a) The absence of PH domain (ΔPH) or coiled-coil-motif-containing tail (ΔCC) does not affect the GAP activity of PpREN. (b) Summary of localization and rescuing ability of mutant forms of PpREN. CDZ, cortical division zone. Cyto, cytoplasmic. (c) Localization of wild-type and mutant forms of PpREN at the CDZ. Arrows indicate the CDZ. Note that only the CC-motif-containing tail is essential for CDZ localization. Scale bars in all panels: 10 µm.

**Supplementary Table 1. Plasmids used in this study.**

| **Plasmid** | **Description** | **Protein Length (aa)** | **Protein MW [kDa]** | **Fusion Protein Length (aa)** | **Fusion Protein MW [kDa]** |
| --- | --- | --- | --- | --- | --- |
| **Pull-down and activity analyses** | | | | | |
| pPY340 | MBP | 1-367 | 40.3 | 417 | 45.6 |
| pPY376 | MBP-Citrine | 2-238 | 26.8 | 634 | 69.9 |
| pPY287 | His-PpROP4-His | 2-180 | 19.6 | 228 | 24.7 |
| pPY341 | GST-PpROP4 | 2-196 | 21.5 | 433 | 48.7 |
| pPY373 | GST-PpROP4(G15V) | 2-196 | 21.5 | 433 | 48.7 |
| pPY374 | GST-PpROP4(D20N) | 2-196 | 21.5 | 433 | 48.7 |
| pPY344 | MBP-PpRopGAP1 | 2-469 | 52.0 | 881 | 96.8 |
| pPY353 | MBP-PpRopGAP1-Nter | 2-193 | 21.1 | 605 | 65.9 |
| pPY348 | MBP-PpRopGAP1-Cter | 194-469 | 31.0 | 689 | 75.6 |
| pPY349 | MBP-PpRopGAP1-GAP | 194-380 | 21.0 | 582 | 63.8 |
| pPY350 | MBP-PpRopGAP1(H165A) | 2-469 | 52.0 | 881 | 96.8 |
| pPY351 | MBP-PpRopGAP1(R239L) | 2-469 | 52.0 | 881 | 96.8 |
| pPY361 | MBP-PpRopGAP1(R134L/R135L) | 2-469 | 52.0 | 881 | 96.7 |
| pPY345 | MBP-PpREN | 2-921 | 100.7 | 1336 | 145.4 |
| pPY362 | MBP-PpREN-ΔPH | 163-921 | 82.7 | 1172 | 127.4 |
| pPY352 | MBP-PpREN-ΔCC | 2-374 | 41.1 | 771 | 84.1 |
| pPY355 | MBP-PpREN(R204L) | 2-921 | 100.7 | 1336 | 145.4 |
| pPY370 | MBP-PpREN(GAP domain) | 163-374 | 23.1 | 603 | 65.7 |
| pPY434 | MBP-AtRopGAP1 | 2-466 | 51.9 | 878 | 96.6 |
| pPY435 | His-AtROP1-His | 2-196 | 21.5 | 250 | 26.9 |
| **Moss transgenesis** | | | | | |
| pPY122 | PpREN knockout - NTC |  |  |  |  |
| pPY123 | mNG-PpREN knock-in |  |  |  |  |
| pPY149 | mNG-PpRopGAP1 knock-in |  |  |  |  |
| pPY150 | mNG-PpRopGAP2 knock-in |  |  |  |  |
| pPY151 | mNG-PpRopGAP3 knock-in |  |  |  |  |
| pPY155 | mNG-PpRopGAP4 knock-in |  |  |  |  |
| pPY152 | mNG-PpRopGAP5 knock-in |  |  |  |  |
| pPY167 | mNG-PpRopGAP6 knock-in |  |  |  |  |
| pPY166 | PpRopGAP1/3/4/6-sg1 - Hyg |  |  |  |  |
| pPY187 | PpRopGAP5-sg1 - Hyg |  |  |  |  |
| pPY194 | PpRopGAP2/4-sg2 - Hyg |  |  |  |  |
| pPY213 | PpRopGAP1-Citrine - Hyg |  |  |  |  |
| pPY216 | PpRopGAP1-mNG - Hyg |  |  |  |  |
| pPY143 | hb7-pEF1α-Cerulean - G418 |  |  |  |  |
| pPY140 | hb7-pEF1α-Cerulean-PpREN - G418 |  |  |  |  |
| pPY204 | hb7-pEF1α-Cerulean-PpRopGAP1 - G418 |  |  |  |  |
| pPY222 | hb7-pEF1α-mNG-PpREN - G418 |  |  |  |  |
| pPY224 | hb7-pEF1α-mNG-PpREN(ΔPH) - G418 |  |  |  |  |
| pPY225 | hb7-pEF1α-mNG-PpREN(R204L) - G418 |  |  |  |  |
| pPY226 | hb7-pEF1α-mNG-PpREN(ΔCC) - G418 |  |  |  |  |
| pPY229 | hb7-pEF1α-mNG-PpRopGAP1 - G418 |  |  |  |  |
| pPY232 | hb7-pEF1α-mNG-PpRopGAP1-Nter - G418 |  |  |  |  |
| pPY233 | hb7-pEF1α-mNG-PpRopGAP1-Cter - G418 |  |  |  |  |
| pPY234 | hb7-pEF1α-mNG-PpRopGAP1(R239L) - G418 |  |  |  |  |
| pPY235 | hb7-pEF1α-mNG-PpRopGAP1(H165A) - G418 |  |  |  |  |
| pPY251 | hb7-pEF1α-mNG-PpRopGAP1(S150-S193) - G418 |  |  |  |  |
| pPY255 | hb7-pEF1α-PpRopGAP1-mNG - G418 |  |  |  |  |
| pPY257 | hb7-pEF1α-PpRopGAP1(fs)-mNG - G418 |  |  |  |  |
| pPY356 | hb7-pEF1α-mNG-PpRopGAP1(Q122-S193) - G418 |  |  |  |  |
| pPY380 | hb7-pEF1α-mNG-CRIB(PpRopGAP1)-GAP(PpREN) - G418 |  |  |  |  |
| pPY389 | hb7-pEF1α-mNG-PpRopGAP1(R134L) - G418 |  |  |  |  |
| pPY394 | hb7-pEF1α-mNG-PpRopGAP1(R135L) - G418 |  |  |  |  |
| pPY412 | PpROP4-swmNG knock-in |  |  |  |  |
| pPY252 | PpRopGEF4-mCherry - G418 knock-in |  |  |  |  |
| **Yeast-two-hybrid** | | | | | |
| pPY274 | BD-PpROP4(C193A) |  |  |  |  |
| pPY275 | BD-PpROP4(G15V)(C193A) |  |  |  |  |
| pPY276 | BD-PpROP4(T20N)(C193A) |  |  |  |  |
| pPY262 | AD-PpRopGAP1 |  |  |  |  |
| pPY267 | AD-PpRopGAP1(Nter) |  |  |  |  |
| pPY268 | AD-PpRopGAP1(CTer) |  |  |  |  |
| pPY269 | AD-PpRopGAP1(H165A) |  |  |  |  |
| pPY270 | AD-PpRopGAP1(R239L) |  |  |  |  |
| pPY293 | AD-PpROPGAP1-Cter(R239L) |  |  |  |  |
| pPY385 | AD-PpRopGAP1(R134L) |  |  |  |  |
| pPY386 | AD-PpRopGAP1(R135L) |  |  |  |  |
| pPY357 | AD-PpRopGAP1(R134/135L) |  |  |  |  |
| pPY400 | AD-PpRopGAP1-Nter(R134L) |  |  |  |  |
| pPY401 | AD-PpRopGAP1-Nter(R135L) |  |  |  |  |
| pPY406 | AD-PpRopGAP1(Q122-S193) |  |  |  |  |
| pPY410 | AD-PpRopGAP(S150-S193) |  |  |  |  |
| pPY414 | AD-PpRopGAP(S150-S193, H165A) |  |  |  |  |
| pPY263 | AD-PpREN |  |  |  |  |
| pPY271 | AD-PpREN(ΔPH) |  |  |  |  |
| pPY272 | AD-PpREN(ΔCC) |  |  |  |  |
| pPY283 | AD-PpREN(130-607) |  |  |  |  |
| pPY478 | AD-PpREN(130-374) |  |  |  |  |
| pPY511 | AD-PpREN(163-374) |  |  |  |  |
