## Supplementary material for "Two subtypes of GTPase-activating proteins coordinate tip growth and cell size regulation in *Physcomitrium patens*": Description of Additional Supplementary Files

[Description of Additional Supplementary Files](https://static-content.springer.com/esm/art%3A10.1038%2Fs41467-023-42236-z/MediaObjects/41467_2023_42236_MOESM4_ESM.pdf)

File name: Supplementary Data 1

Description: **RopGAP sequences for alignment and phylogenic analysis.** Sequence IDs were obtained by domain architecture search for proteins that contain a CRIB motif (IPR000095) and a RhoGAP domain (IPR000198) in the InterPro database. Sequences and taxonomic lineages were retrieved from the UniProt database. Clades that do not contain the conserved pre-CRIB motif are colored in blue.

File name: Supplementary Movie 1

Description: **Dynamic localization of PpRopGAP1-mNG (green) and Lifeact-mCherry (magenta) in tip-growing caulonema cells.** Images were taken at three-minute intervals and were displayed at a rate of 10 frames per second. Note that the actin foci underwent periodical assembly and disassembly.

File name: Supplementary Movie 2

Description: **Dynamic localization of PpRopGAP1-mNG (green) and PpRopGEF4-mCherry (magenta) in tip-growing caulonema cells.** Images were taken at two-minute intervals and displayed at a rate of 10 frames per second.

File name: Supplementary Movie 3

Description: **3D surface intensity plots of PpRopGAP1-mNG and PpRopGEF4-mCherry over time.** The plots were generated from images taken at two-minute intervals and displayed at a rate of 10 frames per second. Note that the intensity signals of PpRopGAP1 (green arrow) and PpRopGEF4 (magenta arrow) fluctuate at the growing apex.

File name: **Supplementary Movie 4**

Description: **Lateral movement of PpRopGAP1-mNG at the apical membrane.** Images were taken at an interval of 0.5 seconds and were displayed at a rate of 10 frames per second.

File name: Supplementary Movie 5

Description: **Dynamic localization of PpROP4-mNG (green) and PpRopGEF4-mCherry (magenta) in the tip cell (right) and pre-branching subapical cell (left).** Note that PpROP4-mNG was enriched at both lateral surfaces in the subapical cell (branching site not specified yet). A transient accumulation occurred at the basal membrane of the tip cell (white arrows). Images were taken at two-minute intervals and displayed at a rate of 10 frames per second.

File name: Supplementary Movie 6

Description: **Dynamic localization of PpRopGAP1-mNG (green) and PpRopGEF4-mCherry (magenta) in the branching subapical cell.** Note that PpRopGAP1-mNG and PpRopGEF4-mCherry were both enriched at the branching site. A transient accumulation occurred at the basal membrane of the tip cell (white arrows). Images were taken at two-minute intervals and displayed at a rate of 10 frames per second.

File name: Supplementary Movie 7

Description: **The division of a subapical cell in the *ropgap, ren* mutant.** Cells were labeled using an actin reporter Lifeact-mCherry. Images were taken at two-minute intervals and displayed at a rate of 10 frames per second.
